# A morphotype of *Saccolaimus saccolaimus* (Chiroptera: Emballonuridae) from West Bengal, India with a comparative note of the species

**DOI:** 10.1101/2023.09.29.560262

**Authors:** Somnath Bhakat

## Abstract

An adult *Saccolaimus saccolaimus* Temminck, 1838 is described here with morphometric measurements in detail. The present species though showed morphological similarities with the species reported from India and abroad but it bears an exceptional character, a white oval pad at the base of the thumb. A comparative note of the species found in India and other countries are also presented here with a suggestion to rethink the IUCN status of the species “Least Concern” in India.

## Introduction

The family Emballonuridae of the order Chiroptera consists of 51 species with a wide distribution in the tropical and subtropical regions of the world e. g. Asia, Africa, Americas, Australia, Europe and Oceania (Mickleburgh et al. 2002). It is commonly known as sheath-tailed bats as the basal part of the tail is enclosed in the interfemoral membrane and the tip protrudes from the upper surface of the membrane and lies free on the dorsal side. Six species of Emballonuridae belonging to two genera, five from the genus *Taphozous* (*T. longimanus* Hardwicke, 1825; *T. melanopogon* Temminck, 1841; *T. nudiventris* Cretzschmar, 1830-31; *T. perforatus* E. Geoffroy, 1818 and *T. theobaldi* Dobson, 1872) and one from the genus *Saccolaimus* (*S. saccolaimus* Temminck, 1838) are recorded from the Indian subcontinent (Bates and Harrison, 1997; Das, 2003; Saikia, 2018). Of these two genera, *Saccolaimus* is one of the biggest emballonurid bats in India, average fore arm length 63.0-68.0mm (Srinivasulu et al., 2010). In India *Saccolaimus saccolaimus* is distributed in Andaman and Nicobar Islands, Gujarat, Karnataka, Kerala, Madhya Pradesh, Maharashtra, Meghalaya, Orissa and West Bengal (Bates and Harrison, 1997; Molur et al., 2002).

These two genera i. e. *Taphozous* and *Saccolaimus* are very difficult to separate externally. Except *T. longimanus* and *S. saccolaimus*, wings are attached to the tibia in other four species of *Taphozous*, while it is attached to the ankles in the aforesaid two species. Moreover, *T. longimanus* bears well developed radio metacarpal pouch which is absent in *S. saccolaimus. Saccolaimus saccolaimus* is the only species of the genus *Saccolaimus* that occurs in South Asia. This is a medium sized bat (FA 63.0-68.0mm), chin with short hairs, second digit without phalanges, conical shaped muzzle without nose-leaf and wings are long and narrow (Srinivasulu et al. 2010). Throat bears a pouch like structure called gular sac which is well developed in males and less developed in female. Tail is of medium size and stout (21.0-35.0mm) (Srinivasulu et al.,2010). Morphology and body measurements of the species are well studied in a few countries like Australia, Bangladesh and Sri Lanka (Troughton, 1925; Chimimba and Kitchener, 1991; Reza et al., 2022; Edirisinghe, 2013). In India, Brosset (1962a) and Dobson (1876) presented body measurements of *S. saccolaimus* in detail. After that, all the studies from different states of India are inadequate. The present paper reports a detailed morphological study of *Saccolaimus saccolaimus* Temminck, 1838 collected from West Bengal, India.

## Materials and methods

The specimen, a mature female was collected from my relative’s house at Suri (87°32’00’’E,23°55’00’’N) Birbhum district, West Bengal, India. The bat was hanging from the ceiling of the third floor of the building.

Measurements of different morphometric features are taken in millimeters. These measurements are taken by using digital calipers or metric rulers. When using a ruler, measurements are taken from the “0” mark (for accurate measurements). Sex was identified by the presence of teat. Weight was measured by a digital single pan balance. Following measurements are taken for the species:

HB (Head and body length): From the tip of the snout to the anus.

FA (Fore arm length): From the extremity of the elbow to the extremity of the carpus with the wings folded from the dorsal side.

EL (Ear length): From the lower border of the external auditory meatus to the tip of the ear.

TL (Tail length): From the tip of the tail to its base to the anus.

HF (Hind foot length): From the extremity of the heel to the tip of the longest digit.

TIB (Tibia length): From the knee joint to the extremity of the heel.

Th (Thumb length): From its base to the nail tip.

2MT (Second metacarpal length): From the extremity of the carpus to the distal end of the second metacarpal.

3MT, 4MT, 5MT (Third, fourth fifth metacarpal length respectively): From the extremity of the carpus to the distal end of the third, fourth and fifth metacarpal respectively.

3D1P, 3D2P, 4D1P, 4D2P, 5D1P, 5D2P (First and second phalanges of third, fourth and fifth digits respectively): From the proximal to the distal extremity of the phalanges.

Tr (Tragus length): From the base of the tragus to its tip.

Ca (Calcar): From the base to its tip.

WS (Wing span): Length between two wing tips.

For comparison all the measurements are presented in percentage of fore-arm length.

## Result

### Taxonomy

Order: Chiroptera Blumenbach,1779

Family: Emballonuridae Gervais, 1855

Subfamily: Taphozoinae Jerdon, 1867

Genus: *Saccolaimus* Temminck, 1838

Scientific name: *Saccolaimus saccolaimus* Temminck, 1838

Synonym: *Taphozous saccolaimus* Blyth, 1844

*Taphozous pulcher* Blyth, 1844

*Taphozous crassus* Blyth, 1844

*Saccolaimus saccolaimus crassus* Blyth, 1844

*Taphozous affinis* Dobson, 1875

*Saccolaimus nudicluniatus* De Vis, 1905

*Taphozous nudicluniatus* De Vis, 1905

*Saccolaimus saccolaimus nudicluniatus* De Vis, 1905

*Saccolaimus pluto* Miller, 1910

Remarks: *S. s. crassus* is considered as a subspecies found only in Indian states. Koopman (1984) and Flannery (1995a, b) considered *S. s. nudicluniatus* to be a subspecies of the widespread *S. saccolaimus*. But Goodwin (1979) questioned the validity of the subspecies status of *S. s. nudicluniatus*.

## Description

Dorsal and ventral view of the species are presented in Fig. 3 and 4 respectively. Morphometric measurements are presented in Table 1. The specimen is large in size (FA 72mm). Proximal phalanges of third finger flexed on dorsal aspect of metacarpal in resting condition. Third metacarpal is longer than the total length of fifth finger including phalanges. Moreover, first and second phalanges of third finger are equal in length. Fifth metacarpal is almost half in length compared to third metacarpal. Toes are almost equal in length. Tragus is less than half to that of ear. The wing membrane is attached to the ankle. Posterior end of the uropatagium is triangular in shape with a pointed tip. Calcar without calcar lobe is present but it is not obvious because of the thickness of the uropatagium. The most important morphometric character of the specimen is the presence of a distinct oval, white thumb pad at the base of the thumb (size of the thumb pad 4.0 x 3.2 mm) (Fig. 1). The species have conical shape muzzle and lack nasal process or nose leaf. Non tubular nostril with tear-drop shaped narial aperture. Lower lip divided by a deep narrow groove in the middle, with a central cushion. Triangular shape pinna with a wide base and have seven ribs on the interior of the pinna. Inner margins of the ear arise from the sides of the fore head and outer margin of the ear conch terminates in a lobe carried forwards towards the angle of the mouth. The distance from the ear end to the angle of the mouth is 5 mm. The distance between the two ear ends on the muzzle is 8 mm. Lower third of the ear is concave, then evenly convex to the tip. The upper three fourth of the outer margin is convex, in lower part of which the ear margin curved inwards to form a thick flattened ledge. In the outer margin of the ear, opposite the tragus, there is a broad semicircular notch which provide the tragus. The medial edge of the pinna smooth and even with the middle of the ear. The tragus has an irregular top edge with three small bumps at one end (Fig. 2). The front surface is concave and hind surface more convex, covered with minute papillae. The tragus has dense hairs on the anterior surface and its base is smooth and even. Tragus is semicircular in cross section. Gular sac represented by a fold of concave integument on the lower and middle one-third portion of the throat (Fig. 1). Metacarpal pouches absent. More than 50% of tail (15.5 mm out of 27 mm) protrudes from the middle of the upper surface of the tail membrane. Tail is not uniform in diameter its base is broad in diameter compared to its end. Distal end of the tail is curved, hook like and its tip bears a few long black hairs.

**Table 1.** Morphometric measurements of *Saccolaimus saccolaimus* from West Bengal (all the values are percentage of FA if not mentioned).

| Parameter | Value | Parameter | Value |
| --- | --- | --- | --- |
| HB (mm) | 91.00 | 3D2P | 38.89 |
| FA (mm) | 72.00 | 4MT | 69.44 |
| EL | 25.42 | 4D1P | 29.17 |
| TL | 37.50 | 4D2P | 9.72 |
| TL (free part) | 21.52 | 5MT | 47.36 |
| HF | 24.30 | 5D1P | 24.03 |
| TIB | 39.03 | 5D2P | 14.58 |
| Th | 6.94 | Tr | 9.58 |
| 2MT | 87.50 | Ca | 31.51 |
| 3MT | 93.33 | WS (mm) | 390 |
| 3D1P | 38.89 | Wt. (g) | 31.32 |

**Fig. 1.**
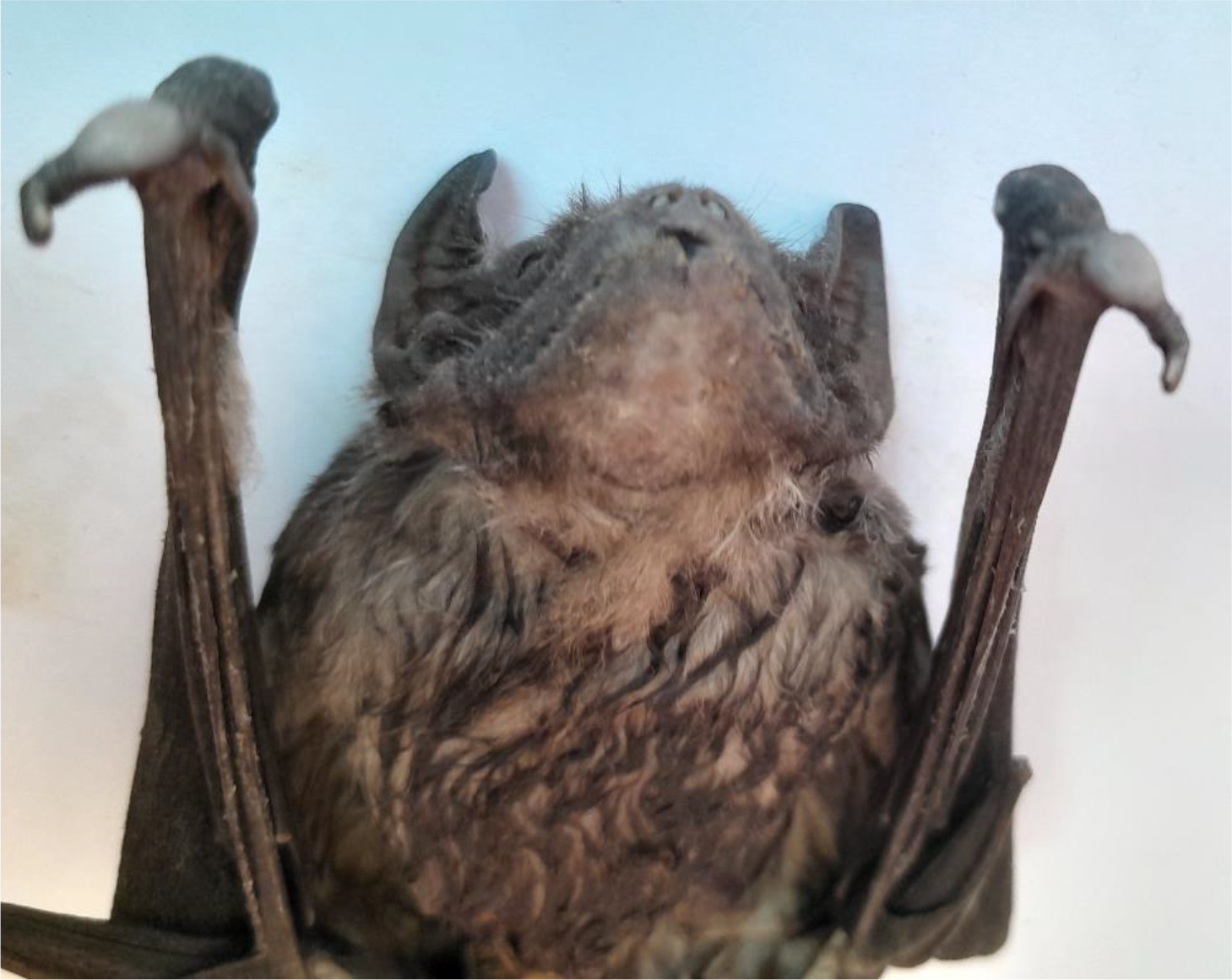
*Saccolaimus saccolaimus*: showing white thumb pad and gular sac.

**Fig. 2.**
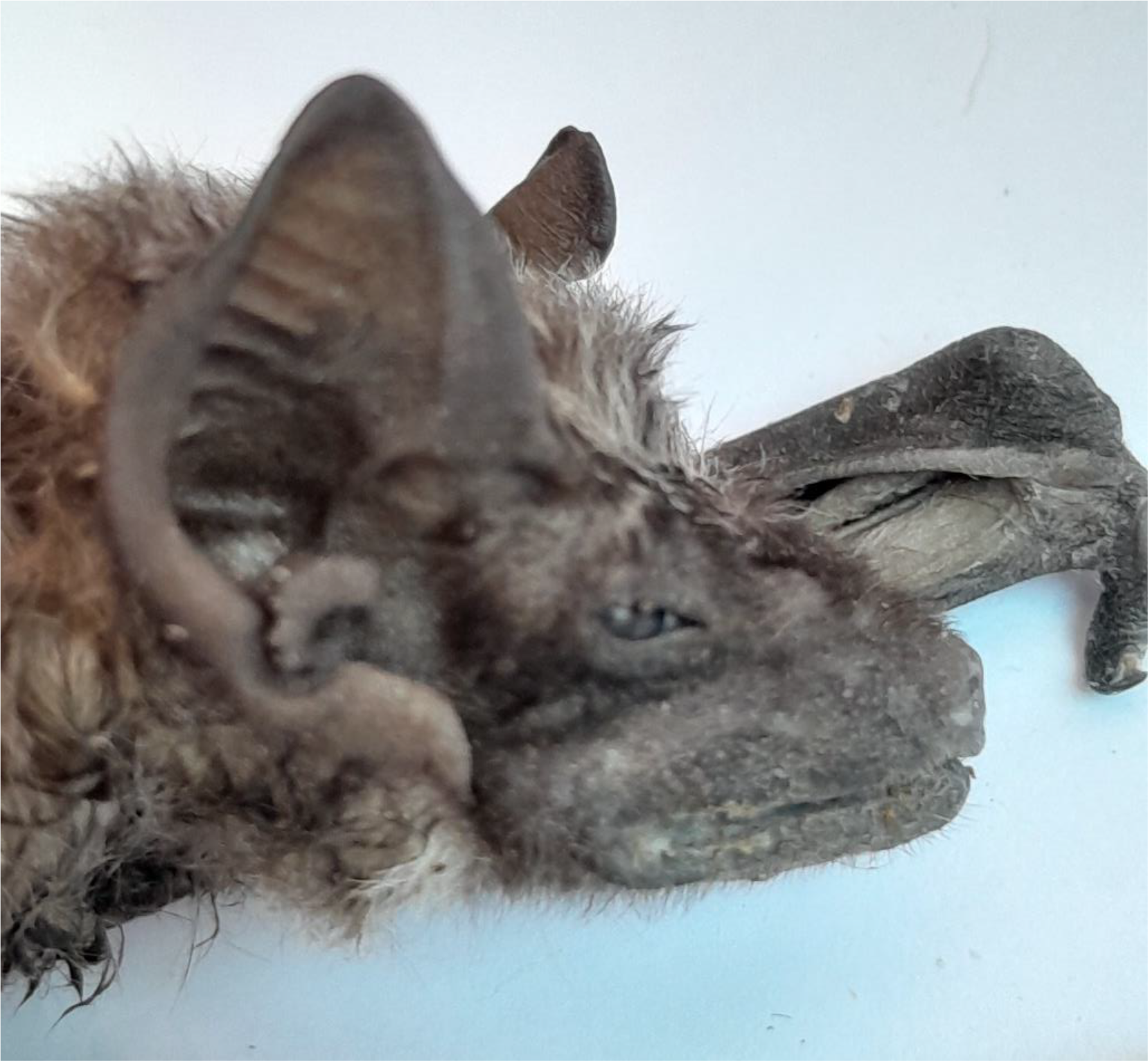
*Saccolaimus saccolaimus*: showing head and pinna with tragus.

**Fig. 3.**
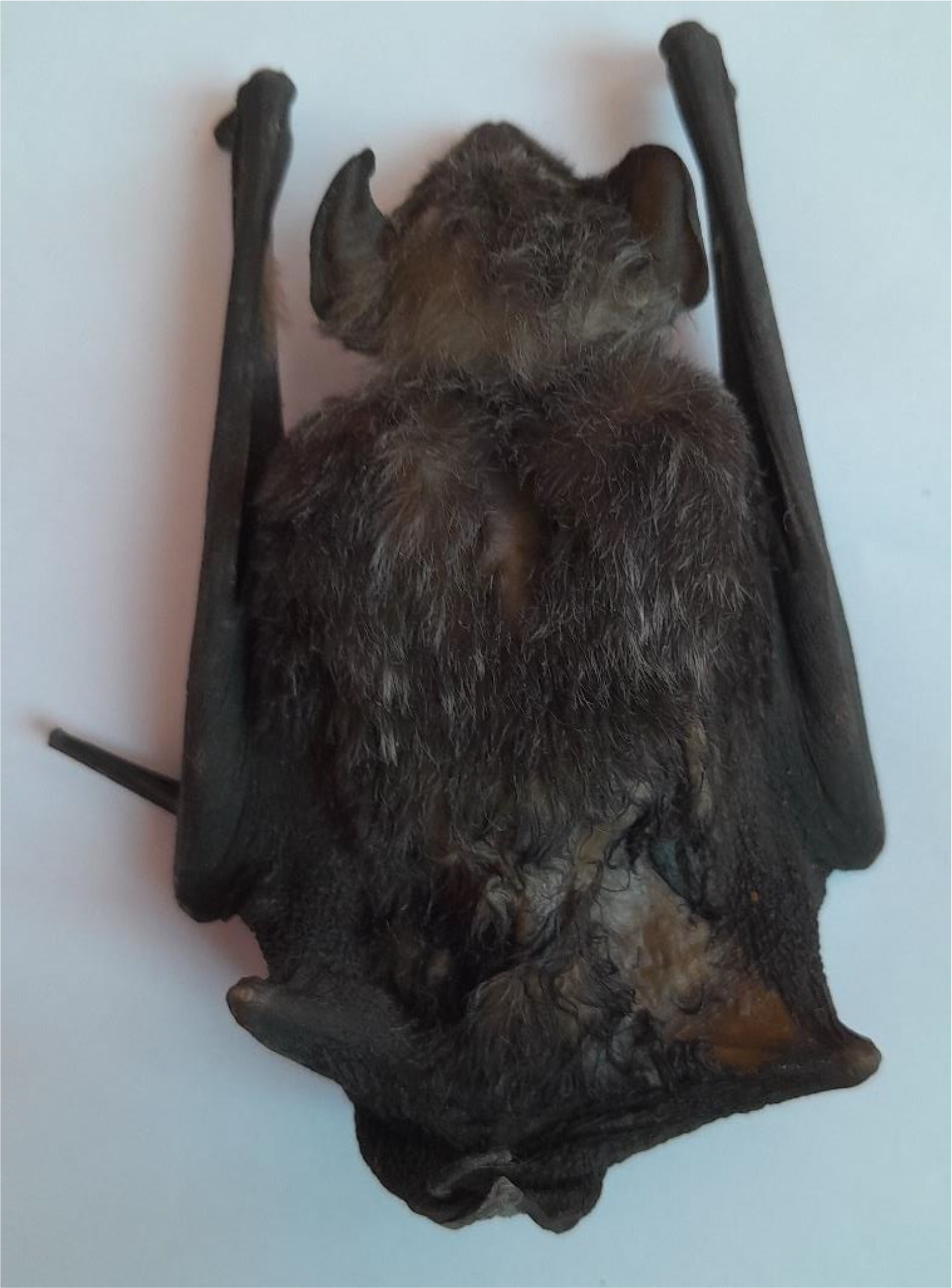
*Saccolaimus saccolaimus*: dorsal view.

**Fig. 4.**
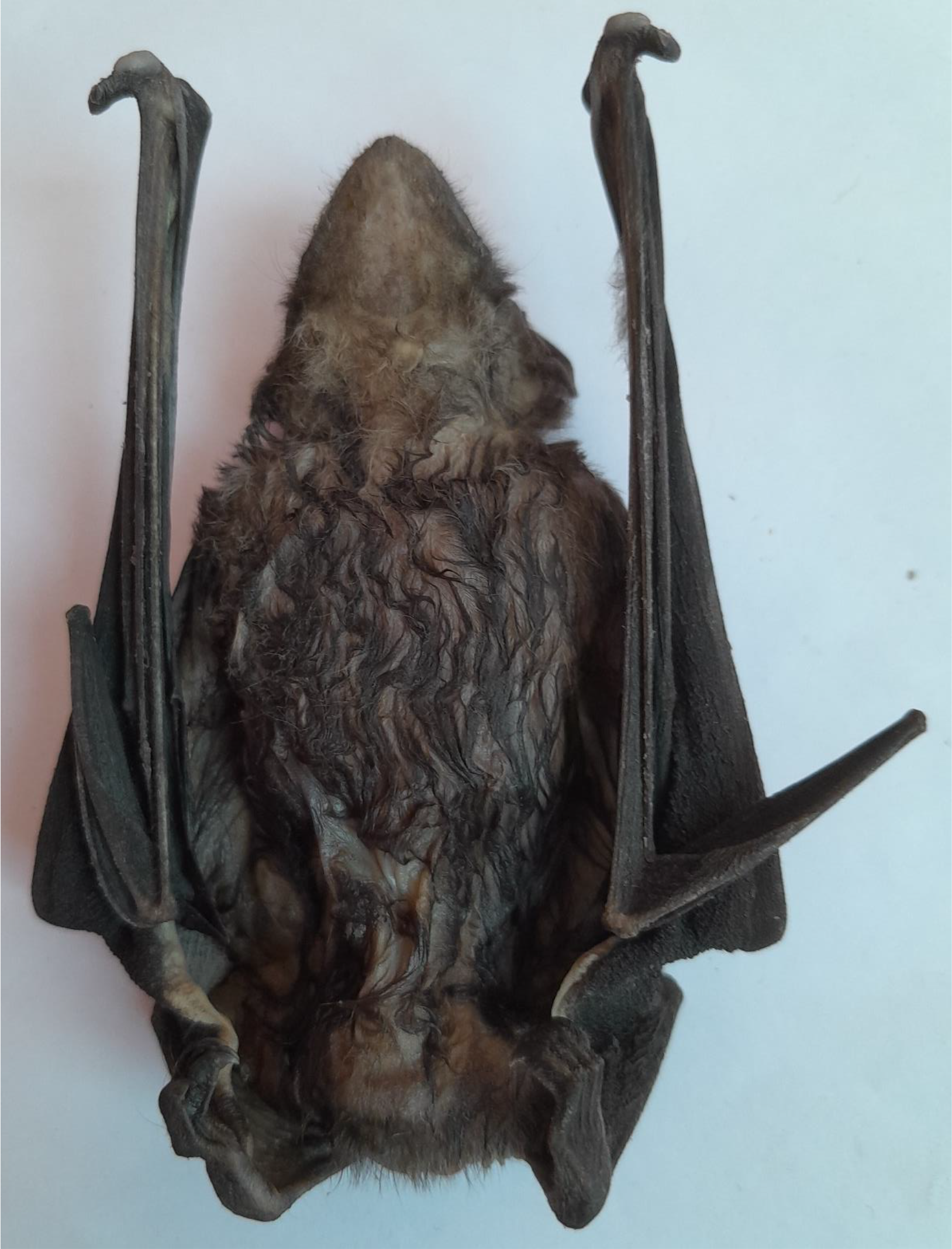
*Saccolaimus saccolaimus*: ventral view.

The dorsal pelage is black with a few small irregular white hairs while ventral pelage less black with white speckle. Interfemoral membrane hairless, thick, spongy and greyish in colour. From fifth finger to the ankle, the margin of the membrane has a white thick border on its dorsal side. Antebrachial membrane with a broad, white margin on its ventral side. Pinna, tibia and patagium without hairs. Face anterior to eye sparsely covered with short black hairs. Tragus with very small hairs. Above on the muzzle hairs are short and scanty. Chin with very fine short hairs except the gular sac encompass. Tail and tibia naked but the tail tip have a few black straight hairs of cf. 5 mm long. Tail is dark black in colour. A tuft of black long straight hairs around the vaginal opening while adjacent hairs are whitish grey. On the dorsal side, long fur extending into the wing to a line between anterior thirds of the humerus and femur. Posteriorly it does not extend past the femur but distributed along a line drawn between their proximal ends. Chest and proximal third of the humerus covered with dark hairs.

A comparative measurement of different external features including weight of *S. saccolaimus* from different countries is presented in Table 2. It indicates paucity of data in different countries except Australia (Troughton, 1925; Chimimba and Kitchener, 1991). Though Reza et al. (2022) from Bangladesh presented a few more measurements of digits. Morphological measurements of *S. saccolaimus* from India are within the range of those reported in the present study. A few characters are uniform in *S. saccolaimus* reported from different countries viz. first and second phalanges of third finger are equal in length, tragus is almost one-third of the ear length, third metacarpal is longer than the fifth metacarpal including phalanges. It is very interesting to note that most of the morphological measurements of the present study are very similar with those reported from Australia though two continents are separated by thousands of miles.

**Table 2.** Comparative study on external morphometric measurements of *S. saccolaimus* in different countries of the world (All values are in percentage of FA if not mentioned). 1. present study; 2. Brossett, 1962a from India; 3. Troughton, 1925 from Australia and New Guinea; 4. Dobson, 1876 from Indian Museum collection; 5. Edirisinghe, 2013 from Sri Lanka; 6. Chimimba and Kitchener, 1991 from Australia; 7. Reza et al., 2022 from Bangladesh; 8. Borissenko and Kruskop, 2003 from Vietnam; 9. Saveng et al., 2011 from Cambodia. (NA-not available).

| Parameter | 1 | 2 | 3 | 4 | 5 | 6 | 7 | 8 | 9 |
| --- | --- | --- | --- | --- | --- | --- | --- | --- | --- |
| HB (mm) | 91.00 | NA | NA | NA | NA | 85.70 | 89.93 | 88.00 | 82.50 |
| FA (mm) | 72.00 | 71.20 | 73.50 | 73.66 | 70.00 | 74.10 | 71.30 | 67.80 | 69.20 |
| EL | 25.42 | NA | 25.85 | 27.59 | NA | 25.24 | 24.56 | 28.61 | 25.14 |
| TL | 37.50 | 39.33 | 35.37 | 44.83 | 34.29 | 43.86 | 38.11 | 35.84 | 41.18 |
| HF | 24.30 | NA | 21.77 | 24.14 | 22.86 | 23.08 | 19.93 | NA | 18.06 |
| TIB | 39.03 | 38.22 | 37.41 | 41.38 | 41.43 | 38.60 | 38.89 | NA | 41.18 |
| Tr | 9.58 | NA | 6.80 | 8.62 | NA | 8.77 | 7.70 | NA | NA |
| 2MT | 87.5 | NA | 89.80 | NA | NA | NA | NA | NA | NA |
| 3MT | 93.33 | 94.12 | 97.96 | 94.83 | NA | 96.63 | 98.20 | NA | NA |
| 3D1P | 38.89 | 44.40 | 40.14 | 39.65 | NA | 37.79 | 39.16 | NA | NA |
| 3D2P | 38.89 | 44.40 | 40.82 | 41.38 | NA | 37.79 | 39.30 | NA | Na |
| 4MT | 69.44 | 71.07 | 70.75 | 51.72 | NA | NA | NA | NA | NA |
| 4D1P | 29.17 | 31.37 | 29.25 | 25.86 | NA | NA | NA | NA | NA |
| 4D2P | 9.72 | 10.96 | 10.20 | 13.79 | NA | NA | NA | NA | NA |
| 5MT | 47.36 | 51.40 | 51.02 | NA | NA | NA | NA | NA | NA |
| 5D1P | 24.03 | 24.72 | 24.49 | NA | NA | NA | NA | NA | NA |
| 5D2P | 14.58 | 15.17 | 16.33 | NA | NA | NA | NA | NA | NA |

Morphometric measurements of six principal characters of *S. saccolaimus* from four different states including the present study are presented in Table 3. It shows that the species from Orissa and Kerala are comparatively smaller in size (FA 64 mm vs. 70 mm) but the other characters are not varied widely except ear length (EL) of Assam species.

**Table 3.** Morphometric measurements of *Saccolaimus saccolaimus* in different states of India (All values are percentage of FA if not mentioned). 1. present study, 2. Das, 2003 from West Bengal; 3. Boro et al., 2013 from Assam; 4. Debata and Palita, 2018 from Orissa; 5. Raman et al., 2021 from Kerala. (NA-Not available).

| Parameter | 1. | 2. | 3. | 4. | 5. |
| --- | --- | --- | --- | --- | --- |
| HB mm | 91.00 | NA | 84.00 | 81.70 | 88.10 |
| FA mm | 72.00 | 69.83 | 70.75 | 64.2 | 64.03 |
| HF | 24.30 | 24.07 | 22.90 | 20.50 | 23.27 |
| TL | 37.50 | 38.85 | 38.16 | 34.73 | 43.70 |
| EL | 25.42 | 28.37 | 21.91 | 27.88 | 28.28 |
| TIB | 39.03 | 39.33 | 39.58 | NA | 36.25 |

## Discussion

The presence of large oval thumb pad at the base of the thumb in the present species is the most peculiar feature which is not reported by any authors earlier. Csorba (personal communication) suggested to study its skull morphology for further confirmation of the species. The differences in the external body measurements between those reported from different countries including South Asia and those presented in the present study is not unexpected as different abiotic factors including types of available food have great impact on the body morphology of a species. In this context, Milne et al. (2002) commented that differences of external characters occurred among individuals are attributed to the same genetic haplogroup and this naturally occurring variations could not be a consequence of subspecific differentiation. Moreover, Chimimba and Kitchener (1991) by PCR analysis showed that phenotypic separation is not evident between *S. saccolaimus* and *S. nudicluniatus* or between the Australian and the Asian *S. saccolaimus*. They also commented that Australian species *S. nuducluniatus* is conspecific with *S. saccolaimus* as supported by Goodwin (1979). Chimimba and Kitchener (1991) and Mckean et al. (1980) observed little variations in *S. saccolaimus* from India to the Solomon Islands. Freeman and Leman (1992) suggested that Emballonurids are less variable in their morphology than are other bat genera. All these reports endorsed the morphological similarity of the present species with those of the non-Indian type especially Australian type.

This species is categorized as ‘Least Concern’ (Lumsden, 2017) both globally and in India (Molur et al., 2002). At present, the distribution of *S. saccolaimus* in different Indian states are not common. Present reports are mainly based on previous record or museum specimen collected earlier. In Madhya Pradesh, it is uncommon and only a single dead specimen was found at Motinala (Ghosh and Bhattacharya, 1995). In Karnataka, no sighting, report based on the study of Bates and Harrison (1997). In Maharashtra, present report is based on old collection of BNHS (Bombay Natural History Society) (Pradhan and Talmale, 2012). In Bihar, Wroughton (1915) reported *T. s. crassus* from Koria, Singbhum district (Singh, 1986). In West Bengal, *S. saccolaimus* was last found in Chandra (Medinipur district in 1984) and Churpurni (Burdwan district in 1988) (Das, 2003). Even in Meghalaya, a state rich in chiropteran fauna, no current report of the species is available (Saikia et al., 2018). *S. saccolaimus* is not reported from a number of states in India e. g. Uttar Pradesh, Rajasthan, Tripura, Manipur etc. Moreover, Saikia (2018) reported the distribution of the species only in four states in India viz. MP, Maharashtra, Assam and Kerala. So, the present IUCN status of *S. saccolaimus*, “Least Concern” in India is questionable. Whereas its IUCN status is ‘Critically Endangered’ in some countries like Australia (Duncan et al., 1999), Sri Lanka (IUCN & MENR, 2007) and ‘Data Deficient’ in Bangladesh (IUCN Bangladesh, 2015).

## Acknowledgement

I am grateful to Mr. Subrata Saha for collection of the specimen. I am also deeply indebted to Dr. Uttam Saikia, Scientist, ZSI for identification of the species and Dr. G. Csorba, Hungarian Natural History Museum, Budapest, Hungary for his comment and suggestion. I am thankful to Dr. P. J. J. Bates of Harrison Zoological Museum. Kent. UK for confirmation of the species.

